# The eyelid and pupil dynamics underlying stress levels in awake mice

**DOI:** 10.1101/2023.08.31.555827

**Authors:** Hang Zeng

## Abstract

Stress is a natural response of the body to perceived threats, and it can have both positive and negative effects on brain hemodynamics. Stress-induced changes in pupil and eyelid size/shape have been used as a biomarker in several fMRI studies. However, there were limited knowledges regarding changes in behavior of pupil and eyelid dynamics, particularly on animal models. In the present study, the pupil and eyelid dynamics were carefully investigated and characterized in a newly developed awake rodent fMRI protocol. Leveraging deep learning techniques, the mouse pupil and eyelid diameters were extracted and analyzed during different training and imaging phases in the present project. Our findings demonstrate a consistent downwards trend in pupil and eyelid dynamics under a meticulously designed training protocol, suggesting that the behaviors of the pupil and eyelid can be served as reliable indicators of stress levels and motion artifacts in awake fMRI studies. The current recording platform not only enables the facilitation of awake animal MRI studies but also highlights its potential applications to numerous other research areas, owing to the non-invasive nature and straightforward implementation.

## Introduction

The awake mouse model, under normal or pathological conditions, has been widely used in functional magnetic resonance imaging (fMRI) as a platform for investigating various physiological activities, e.g., brain fluctuations[1–3], behavior[4], cognition processing[5, 6]. Thus, laying a solid foundation for translational studies from basic brain research to the clinical applications. Nowadays, numerous neuroimaging laboratories are working on developing awake animal models for their fMRI protocols. However, one significant challenge needs to be carefully considered, i.e., the potential stress caused to the animals during the imaging procedures, which can severely affect physiological states and brain hemodynamic activities. In previous studies, researchers have utilized various methods to measure stress levels in rodents, such as plasma corticosterone levels[2], changes in blood pressure, respiration rate or heat rate[7–9], etc. Though those measurements can be applied, it is still crucial to consider the limitations and potential confounding factors of each method when applying to the awake animals, particularly for awake fMRI protocols. As a results, many labs are exploring alternative approaches, such as pupillometry and eyelid dynamics, which are non-invasive and effective methods for stress level monitoring [3, 10].

Previous studies have demonstrated the relationship between stress and pupil dynamic[11]. One study found that pupils of individuals exposed to stress-inducing stimuli, such as a loud noise or electric shock, dilated more than pupils exposed to non-stressful stimuli[12]. Another study found that the pupils of individuals with post-traumatic stress disorder (PTSD) were more likely to remain dilated than those without PTSD, indicating a persistent stress response[13]. In addition to pupillometry dynamics, eyelid dynamics have also been proposed as a promising biomarker of stress in both humans and animals. For example, one study found that individuals with high levels of anxiety and stress exhibited increased blink rates and reduced eyelid closure duration[14]. Previous studies have reported eyelid behaviors, i.e., closure, open, and blinking, etc., have been linked with brain fluctuations[15–18]. The eyelid closures were reported be to link with the brain arousal state[15]. Whereas eye-blinking events were found related to cortical fluctuations[18]. Furthermore, the eyelid closures and openings modulate brain functional connectivity, showing different temporal properties of brain networks[19].

The pupil and eyelid behaviors conveying massive information flows, have been applied as the physiological tenants to underlie the stress levels. Nonetheless, the significant pupillometry dynamics and eyelid behavioral states lack a precise evaluation for encoding stress levels. Here, in the current project, we aim to non-invasively investigate the possible link between MRI-induced stress levels and pupil/eyelid dynamics on an awake mouse model, including how the pupil/eyelid changes in size and shape occur in response to stress, and how these changes, particularly, the the dynamic temporal features, can be utilized to evaluate the stress levels in building the awake fMRI protocols. By comprehensively characterizing the pupil/eyelid dynamics and their relationship with stress levels, we seek to provide a novel and reliable approach for assessing stress in awake animal imaging modalities, and establish the validity and reliability of this approach with potential applications in both preclinical research and clinical practice.

## Material and methods

### Animals and habituation

7 male C57BL/6 mice (Charles River Laboratory) were trained in this project. Animals were habituated individually at a 12 hr-12 hr light-dark cycle (light on from 8 a.m. to 8 p.m.) with food and water ad libitum, starting at age 10-15 weeks. All the surgical and imaging procedures were approved by the state authority (Regierungspräsidium, Tübingen, Baden-Württemberg, Germany) and conducted in full accordance with the German Animal Welfare Act (TierSchG) and Animal Welfare Laboratory Animal Ordinance (TierSchVersV).

All the animals were trained 5-8 weeks for the awake MR imaging. The animals’ head was fixed, and the body left unrestrained within the training apparatus. Only the well-trained animals were applied in further imaging. The acclimating procedures were depicted in our previous report[20].

### Head-post surgery

Mice were anesthetized by isoflurane (2% for induction; 1-1.3% for surgery in a mixture of 30% oxygen in air) and fixed in a stereotaxic frame. The customized head-post was printed out using our 3D printer(From 2, www.formlabs.com). We applied dental cement (Charisma flow) to fix the head-post on top of the skull. Painkillers and antibiotics were administered to relieve pain and inflammation immediately after finishing the surgery. Animals were maintained on a heated blanket until fully recovered from the anesthetic effect (15-30 mins) and were placed back in their original cages. Post-surgical observations were documented for a minimum of five consecutive days following the surgery (twice a day for five days and then daily). Detailed surgical procedures can be found in our previous reports.

### Eyelid/pupil acquisition

A customized MRI-compatible camera with infra light (850 nm) was specially designed and applied for eyelid tracking and pupillometry recording in this study. Under iso-/luminance, the pupil and eyelid videos (32 bites/pixel, with a resolution of 1920×1080 pixel and refresh rate of 60 frames/s) were continuously recorded from left eye of the mouse.

The camera was mounted in an adjustable non-magnetic holder for consistence of MRI scanner. The animals were head-fixed and positioned ∼0.8 cm away from the camera in the restraining apparatus. The eyelid/pupil video recordings were started simultaneously with the initial of MR imaging.

### fMRI data acquisition

The fMRI datasets were acquired using a Bruker Avance III System (Bruker BioSpin, Ettlingen, Germany) with a 14.1-T superconducting magnet with a 12 cm diameter gradient providing 100 G/cm with a 150 µs rise time. A custom-made transceiver surface coil (8 mm in diameter) covering the whole brain of mice was applied for imaging. Functional scans and anatomical images were acquired using the 2D echo-planar imaging (EPI, 24 slices, 0.3×0.3×0.5 mm^3^resolution) and rapid acquisition with relaxation enhancement sequences (RARE, 24 slices, 0.125 mm^2^ in-plane resolution, 0.5 mm slice thickness) sequences, respectively. All the fMRI datasets were preprocessed by using the Analysis of Functional NeruoImage (AFNI) software package (https://afni.nimh.nih.gov/)[21].

### Data analysis

The coordinates of eyelid/pupil were automatically labeled and extracted using the DeepLabCut [22, 23] network. To establish this network, 37 raw videos were applied and trained in the current study. Basically, 100 frames/video were chosen and labeled for training the network (8 points per frames, i.e., right, left, up, bottom boundary of eyelid and pupil, respectively). The algorithm for measuring the diameter of eyelid (ED) and pupil(PD) is as following[3, 24]:

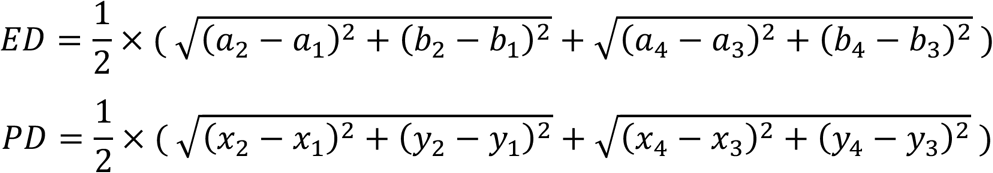

Where the coordinates of the points (*ai, bi*), *i* = 1, …, 4 and (*xj, yj*), *j* = 1, …, 4, were used to calculate the ED and PD, respectively.

The anatomical MR image qualities were assessed by comparing the SNR (signal-to-noise-ratio) between the different time points extracted from the training datasets. The formular of SNR is as following[25]:

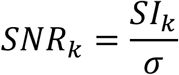

where 𝑆𝐼*k* stands for mean of signal intensity in the regions of interest (ROIs) labeled by *k*, and 𝜎 denotes the standard deviation of background signal intensity outside the ROI. We manually selected the corpus callosum (close to the surface and shows less distortion) as the ROI to represent the whole brain image and calculate the signal intensity.

### Statistical analysis

All data are shown as mean ± SEM. Statistical analyses were performed in python or GraphPad Prism version 9, and all figures were generated in python and GraphPad.

## Results

### Experimental design

Figure 1 shows the schematics of the awake mouse eyelid and pupillometry recordings, as well as the training system and MR imaging modules. Figure 1A depicts the experimental designs of the animal, undergoing headpost implantation, restraint training, and fMRI imaging, etc. The experimental procedures matching to the training and imaging steps were shown in Fig.1B, e.g., a plastic headpost was firstly, implanted and then trained in the customized apparatus, and finally, went for imaging in MR scanners. Fig.1C shows the eyelid and pupil diameter were recorded and extracted for analysis. The functional and anatomical MR images were also obtained when the animals were trained ready for imaging.

**Figure 1.**
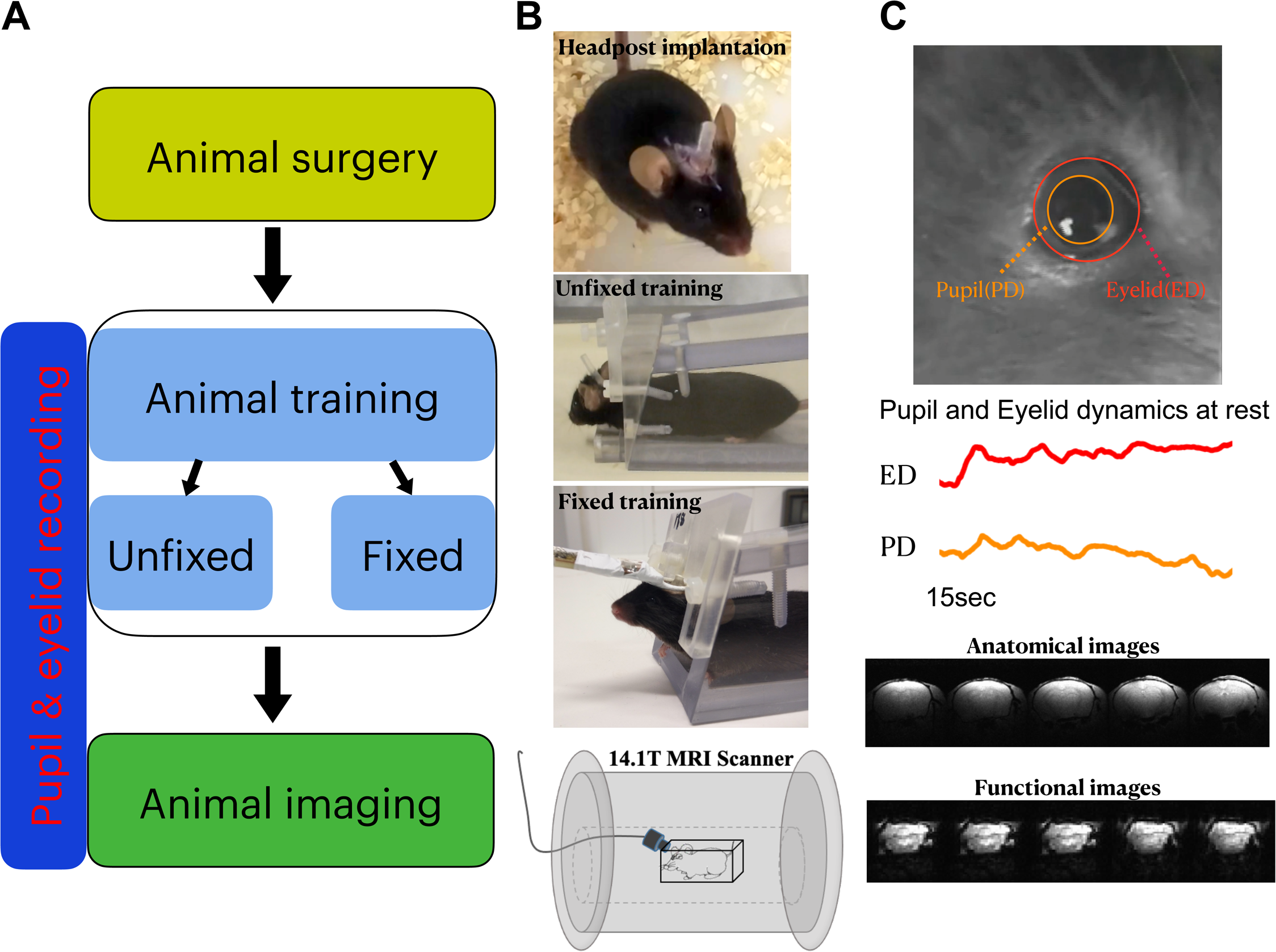
The work flow of the experimental designs. Fig.1A depicts the experimental designs of the animal, undergoing headpost implantation, training, and MRI imaging. Fig.1B shows the experimental demos matching to the experimental techniques in Fig.1A. Fig.1C is a representative trial of the eyelid and pupil recordings and MR images.

### Pupil dynamics

Figure 2 shows the pupillometry dynamics during the training period. Of which, Fig.2A displays the mean pupil diameter from all the recorded trails in different training sections. Moreover, the pupil diameter correlated with the training period, showing a decline trend towards the end of restraining section (linear regression, R^2^=0.55, *p*<0.01). Further analysis revealed significant changes of the pupillometry following the implemented training protocol, i.e., earlier phases (1-3 week) vs. later phases (4 - 11 week), showing reduced pupillometry dynamics (Fig.2B, p<0.01). These results may suggest that the animals get accustomed to the training and show less stress to the restraint apparatus. It thus, further validates the current awake mouse MRI platform for brain imaging.

**Figure 2.**
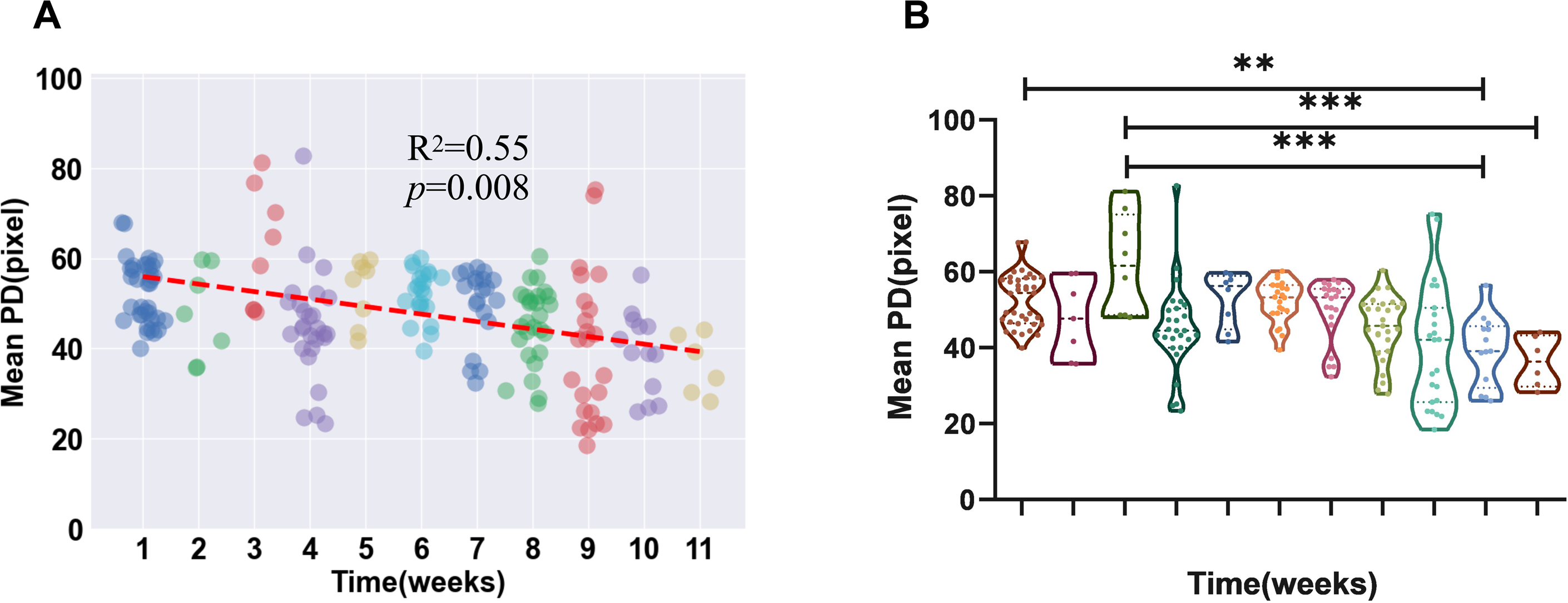
Pupil dynamic changes following the training weeks. Fig.2A shows the scatter plot of the mean pupil dynamics and the linear regression of those plots following the training periods. Red line denotes the trend of the changes for the pupil. Fig.2B shows the statistical analysis of the mean pupil diameters among the different weeks (mean±S.E.M, one-way ANOVA, ***p<0.001, Bonferroni post hoc test).

### Eyelid dynamics

Similarly, the eyelid dynamics from every individual animal were extracted through our trained DeepLab Cut network. The average eyelid diameter across all training sections were graphically represented as a function of time, as illustrated in Fig.3A, which revealed that later training sections had fewer alterations. Moreover, a linear regression analysis was also conducted to assess the changes in eyelid dynamics, which yielded a coefficient of determination (R^2^) value of 0.64 (p<0.01), indicating a significant decline in eyelid dynamics. Further statistical analysis verified the identical findings of a smaller eyelid dynamics in the late stage (5-11 week) of training in comparison to the early stage (1-4 week) of training (Fig.3B, p<0.01). These findings imply that the animals, after intensive training in the MR apparatus, became habituated to the imaging environment and showed decreased discomfort.

**Figure 3.**
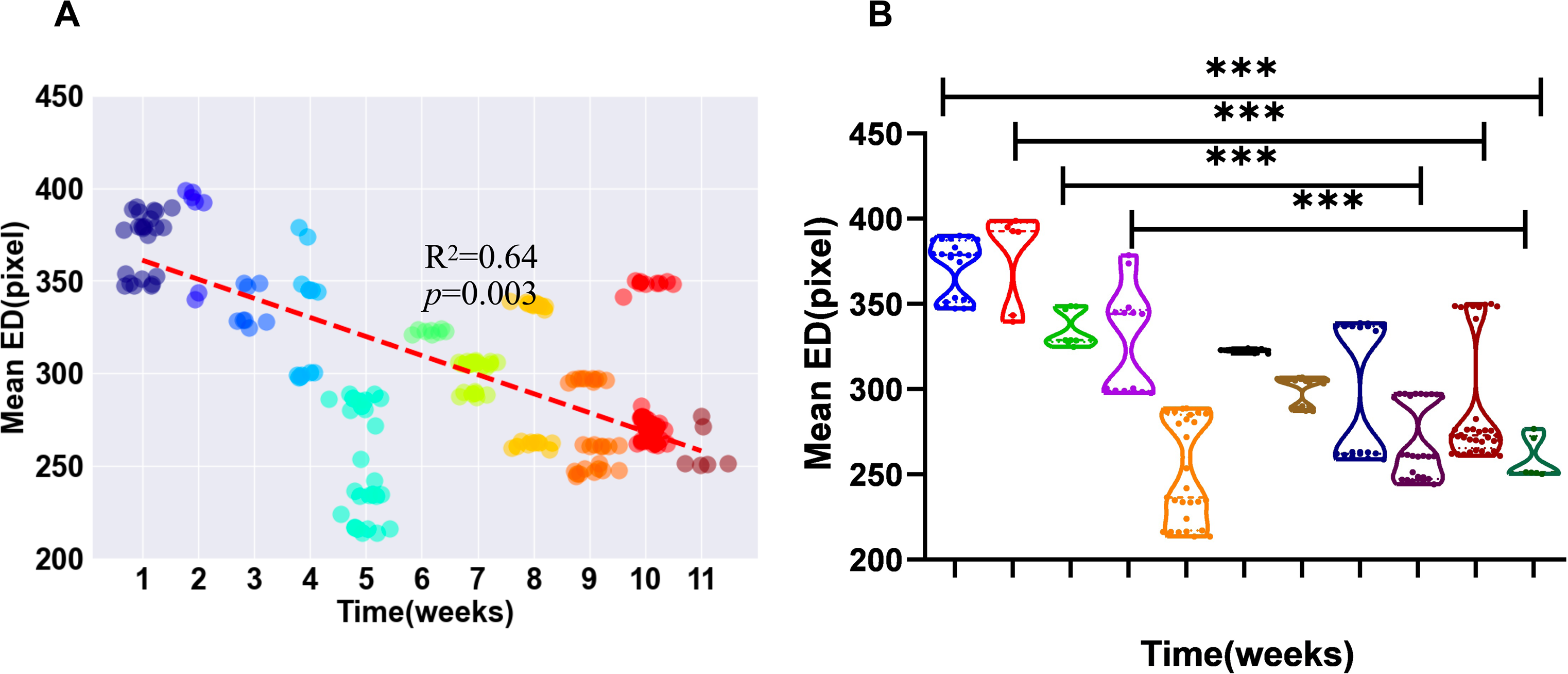
Eyelid dynamic changes following the training weeks. Fig.3A shows the scatter plot of the mean eyelid dynamics and the linear regression of those plots following differnent training periods. Red line denotes the trend of the changes for the eyelid diameters. Fig.3B shows the statistical analysis of the mean eyelid diameters among the different weeks (mean±S.E.M, one-way ANOVA, ***p<0.001, Bonferroni post hoc test).

### SNR analysis of the MR images

Next the MR image quality were also quantified by calculating the SNRs from different training stages when the brain images were acquired. As depicted in Fig.4A, the SNR values pertaining to the spatial MR images exhibited a gradual improvement throughout the training periods, indicative of a progressive upward trend towards the training phase (R^2^=0.71, p<0.05). Further analysis suggested that the SNR of the MR images acquired were better in the late stage (7-11 week) than early stage (1-6 week) (Fig.4B, p<0.05). The observed augmentation in SNR for the MR images implies a reduction in motion artifacts during the image acquisition process, thus signifying a diminished level of stress and discomfort of the animals.

**Figure 4.**
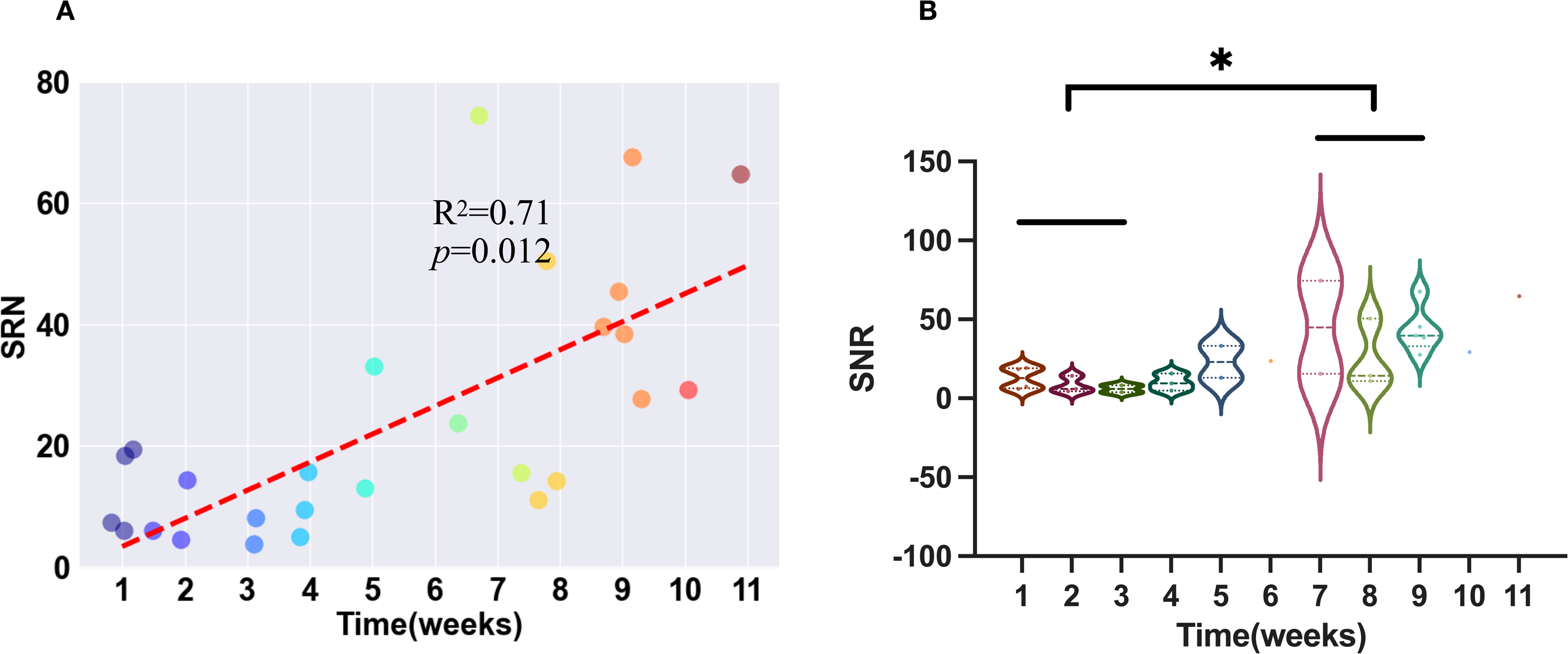
The SNR of the MRI scans among the different weeks of training. Fig.4A shows the scatter plot of the mean SNR values and the linear regression of those plots at different weeks of training. Red line denotes the trend of changes for SNR. Fig.4B shows the statistical analysis of the SNR values among the different weeks (mean±S.E.M, one-way ANOVA, *p<0.05, Bonferroni post hoc test).

## Discussion

Pupillometry, a technique that measures the pupil’s size and dynamics to assess the cognitive, mental demands and arousing state levels, etc., in individuals that has long been investigated for over 50 years[26]. Nowadays, it has led to the development of various methods for measuring stress levels for both human subjects and animal models. The current study, by applying the deep neural networks, not only presents the dynamics changes of pupillometry, but also shed light on the dynamics of eyelid for monitoring the physical stress on an awake mouse model, particularly, during the fMRI imaging modalities. Making it a valuable tool to non-invasively assess the stress levels and cognitive overload in individuals with potential applications for future preclinical researh and clinical practice.

The animals in the current study were consciously being exposed to such an acoustic environment (>100 dB, produced by the scanner) and were kept still without movements. It is thus impossible for the animals to avoid the physical stress induced by the imaging apparatus. To mimic the stress-induced changes in brain hemodynamics, we have applied a new awake training protocol and kept measuring the pupil diameter changes throughout the entire experiment. Our findings proved that with proper training procedures and exposure periods, the animals’ training effects can be visualized and quantified as indicated by the pupillometry measurement in a real-time manner. Instead of measuring corticosterone level or recording respiration/heart rate as the stress biomarkers, we assessed the dynamic changes of the pupil due to its non-invasive and non-physical contact to the conscious animals. Furthermore, stress-induced changes in pupil size and shape have been suggested to be resulted from the activation of the autonomic nervous system’s sympathetic branch[27, 28]. Therefore, the pupillometry, on the one hand reflects the stress levels during training, and on the other hand, indicates the activities of the sympathetic nervous system while imaging. Ensuring the examiners to perform longitudinal studies and to explore the brain hemodynamics underlying behavior or disease states with limited stress-induced effects in a controlled laboratory setting.

The use of eyelid dynamics to assess the stress-induced effects has been reported in previous studies[29]. However, very few studies have focused on the eyelid dynamics and the examination of eyelid dynamics as a potential stress biomarker remains largely unexplored. Though, preliminary findings suggest that increased blink rates[30], shorter durations of blinks[15], and enhanced eyelid closure velocity[15, 31] were all related to higher levels of stress. However, these results underestimated the potential applications of eyelid dynamics as a promising stress biomarker. In the current investigations, we dynamically retrieved and analysed the changes in eyelid diameter on a mouse model, revealing a substantial link between stress levels and changes in eyelid dynamics, particularly during training and imaging for fMRI modalities. Our analysis indicated that the eyelid dynamics provide significant valuable insights between the earlier and later phases of training/imaging modules, offering a non-invasive and objective insights into the stress-induced response on awake animal models.

In conclusion, this study sheds light on the relationships between stress level and pupil & eyelid dynamics, highlighting the potential applications of pupil and eyelid dynamics as a non-invasive and objective stress biomarker. By monitoring the pupil and eyelid diameter changes, researchers and clinicians can potentially identify and measure stress levels in real-time, enhancing the accuracy and efficiency of stress assessment. Our findings may also provide a step toward understanding of the complex interplay between stress and human behavior, which might revolutionize stress assessment and contribute to the development of personalized stress management strategies in various domains, including healthcare, occupational settings, and mental well-being, etc.

### Limitations of the study

Overall, measuring stress levels in awake animals is essential, not only for ensuring well-being of the animals, but also critical for investigators to produce reliable and repeatable results. We have proposed to measure the dynamics of the pupil and eyelid as the biomarkers for stress evaluations. However, there are several limitations that must be considered. One of the main limitations is that the pupil and eyelid dynamic data were recorded in a controlled, specific fMRI lab settings. Such a constraint and noisy environment may not only induce physical stress to the conscious animals, but also might be psychological-associated as well. It may vary depending on the examiners’ experimental surroundings. Another limitation is the lack of standardization between the stress levels and the dynamics of pupil and eyelid. By real-time monitoring the diameter changes of pupil and eyelid, the stress-induced effects can be detected. However, to what extent this effect can be indicative and interpretable remains ambiguous, particularly on awake animal models. Finally, the pupil and eyelid are not a perfect circle, and its shape and position can be affected by changes in the iris’s curvature and illuminations. Future investigations may focus more on developing new analytical methods as well as the implementation of well-designed experiments.

## Acknowledgements

We thank Miss. Gao for processing the raw videos and providing analytical comments.

## Author contributions

Dr. Zeng and Miss. Gao were responsible for the data analysis as well as the preparation of the manuscripts.

---

**Supplementary video S1.**
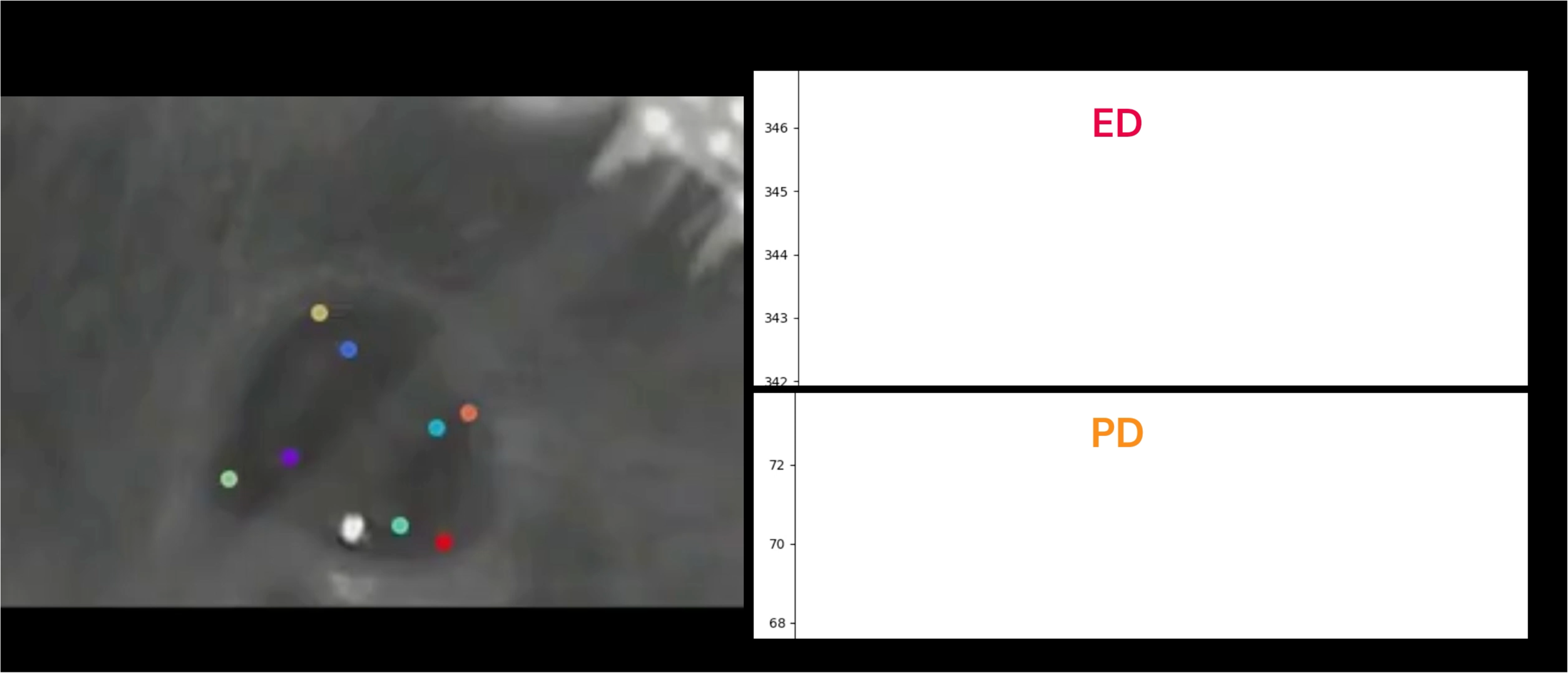
The left video shows the labeled pupil and eyelid dynamics, and the right video shows the corresponding timecourses of eyelid dynamics (ED) and pupil dynamics (PD).

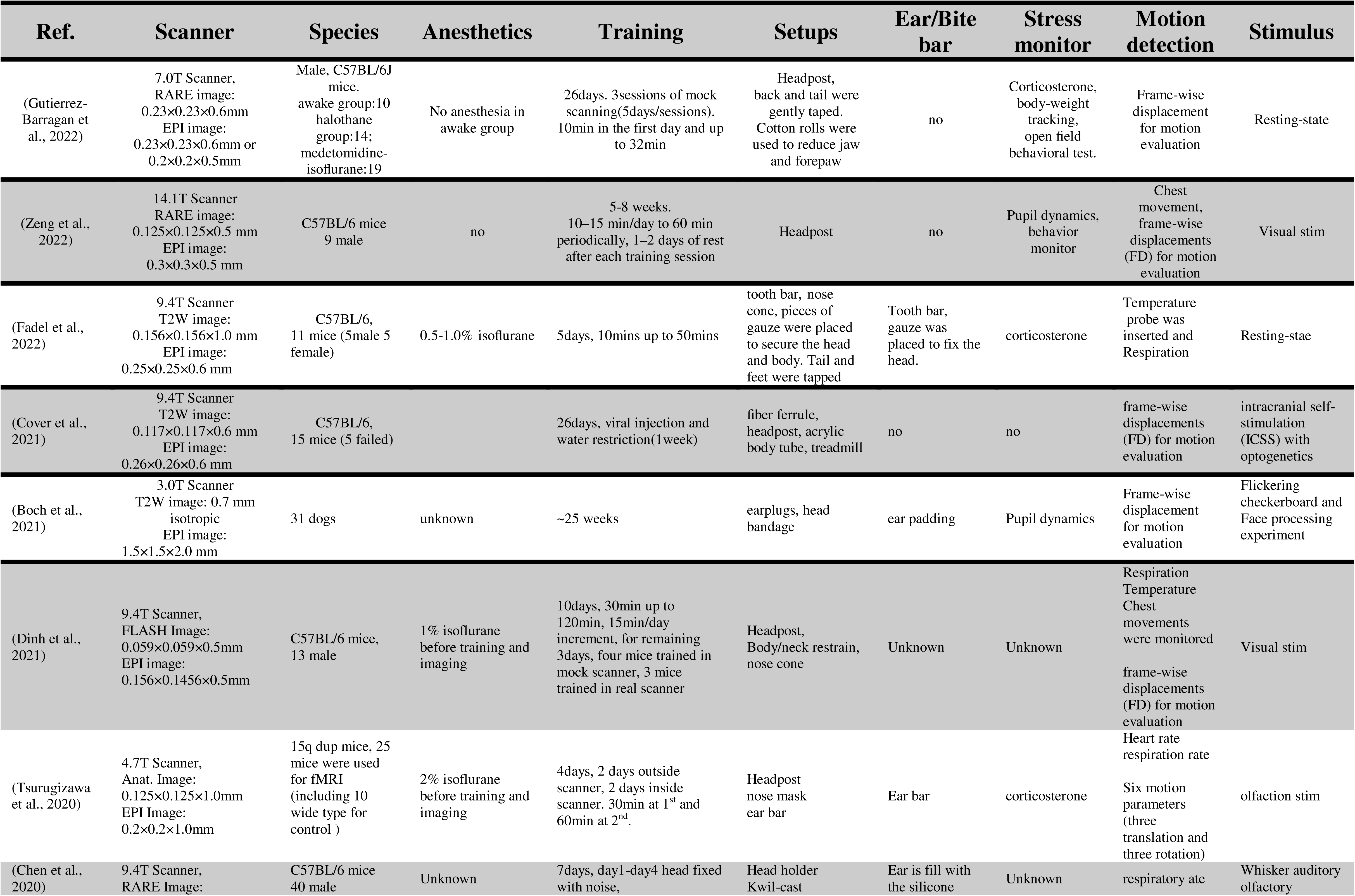
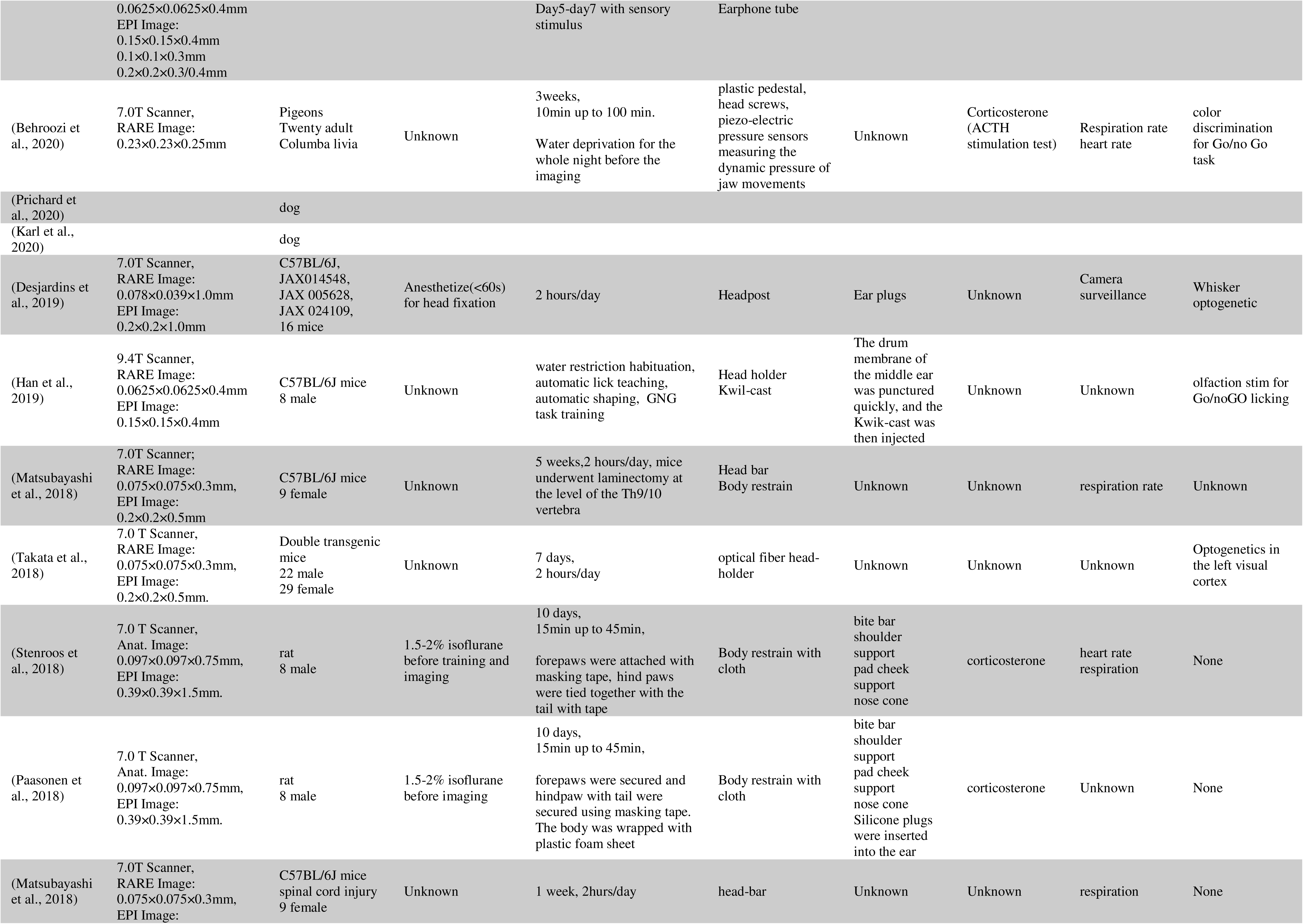
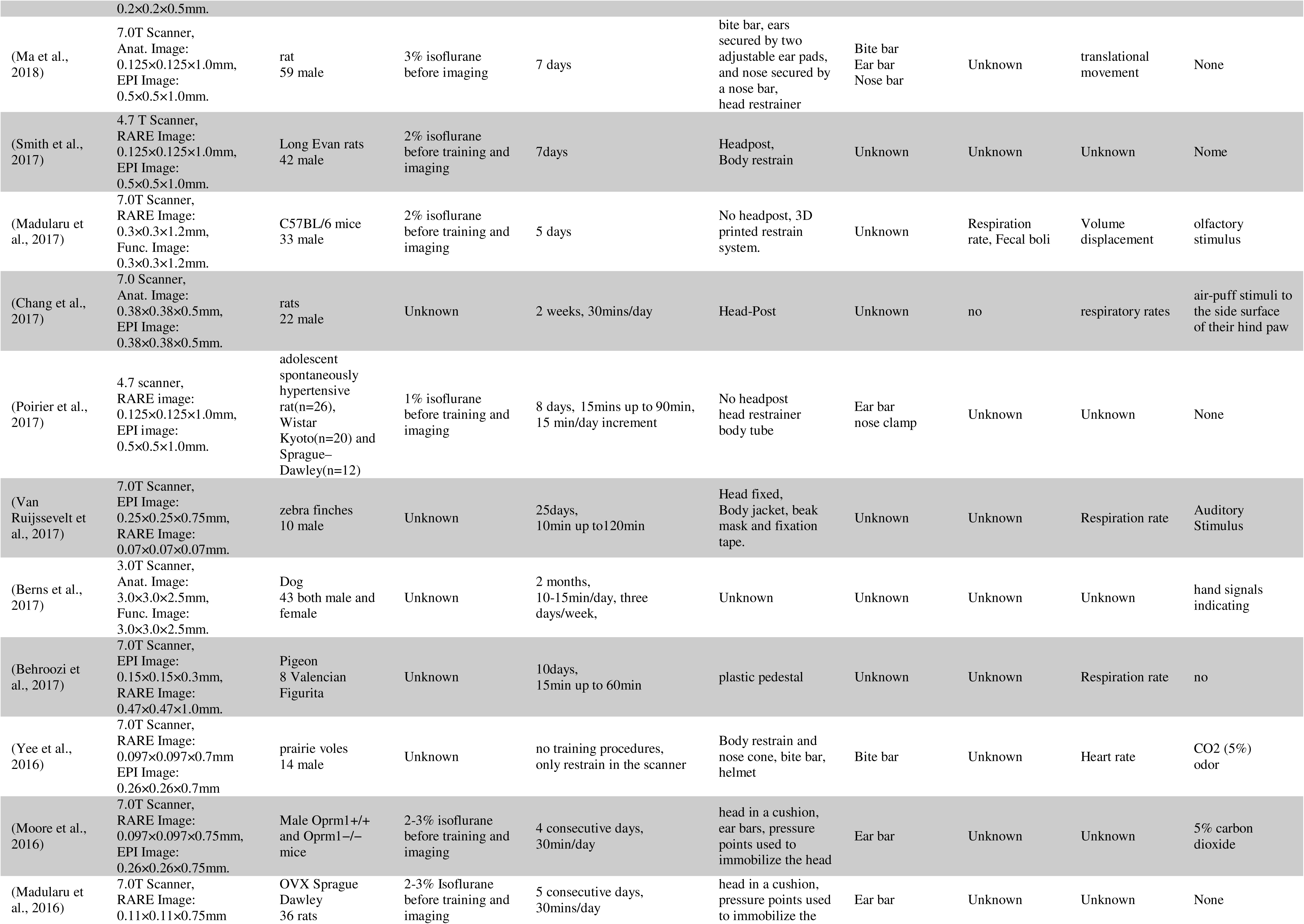
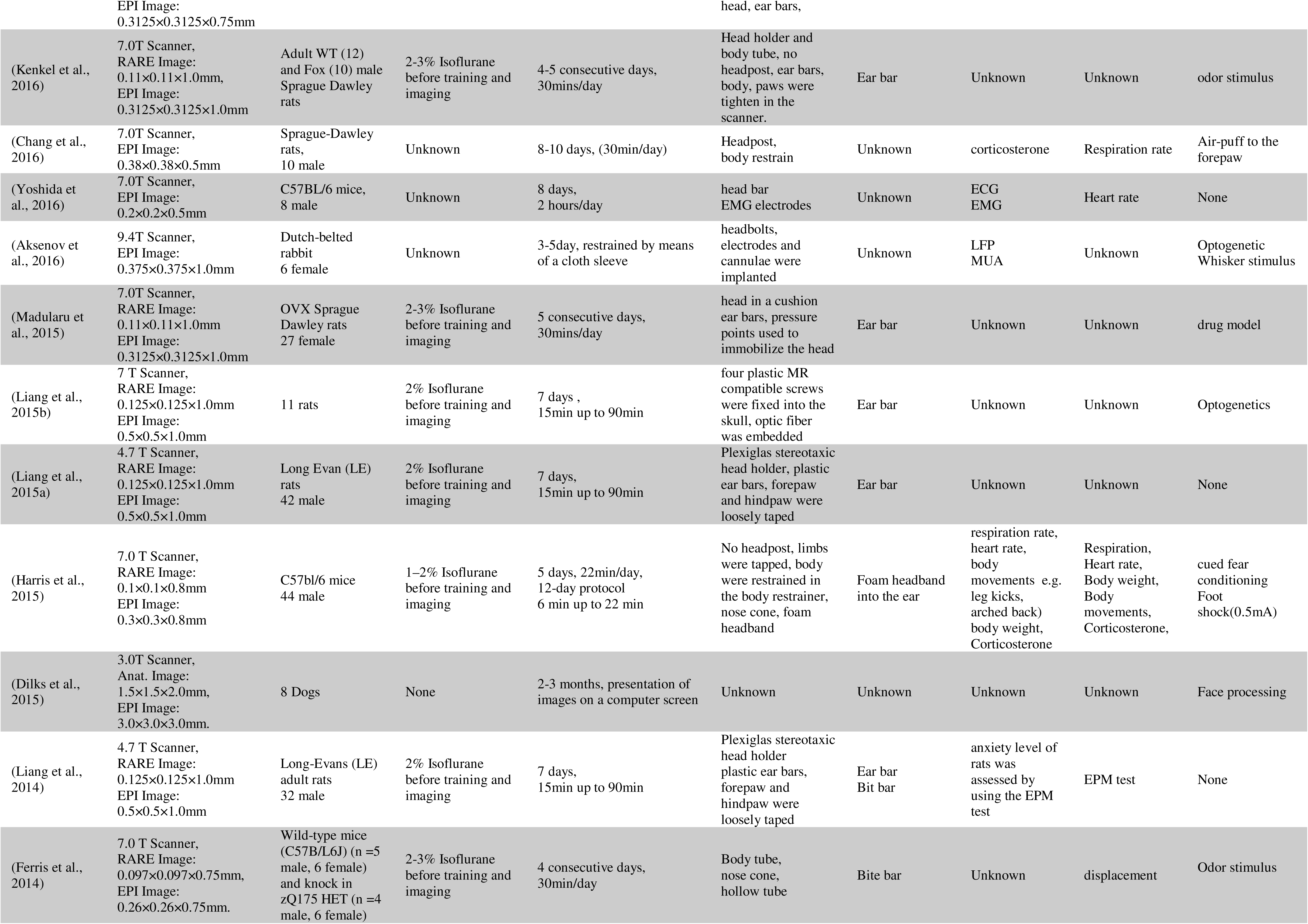
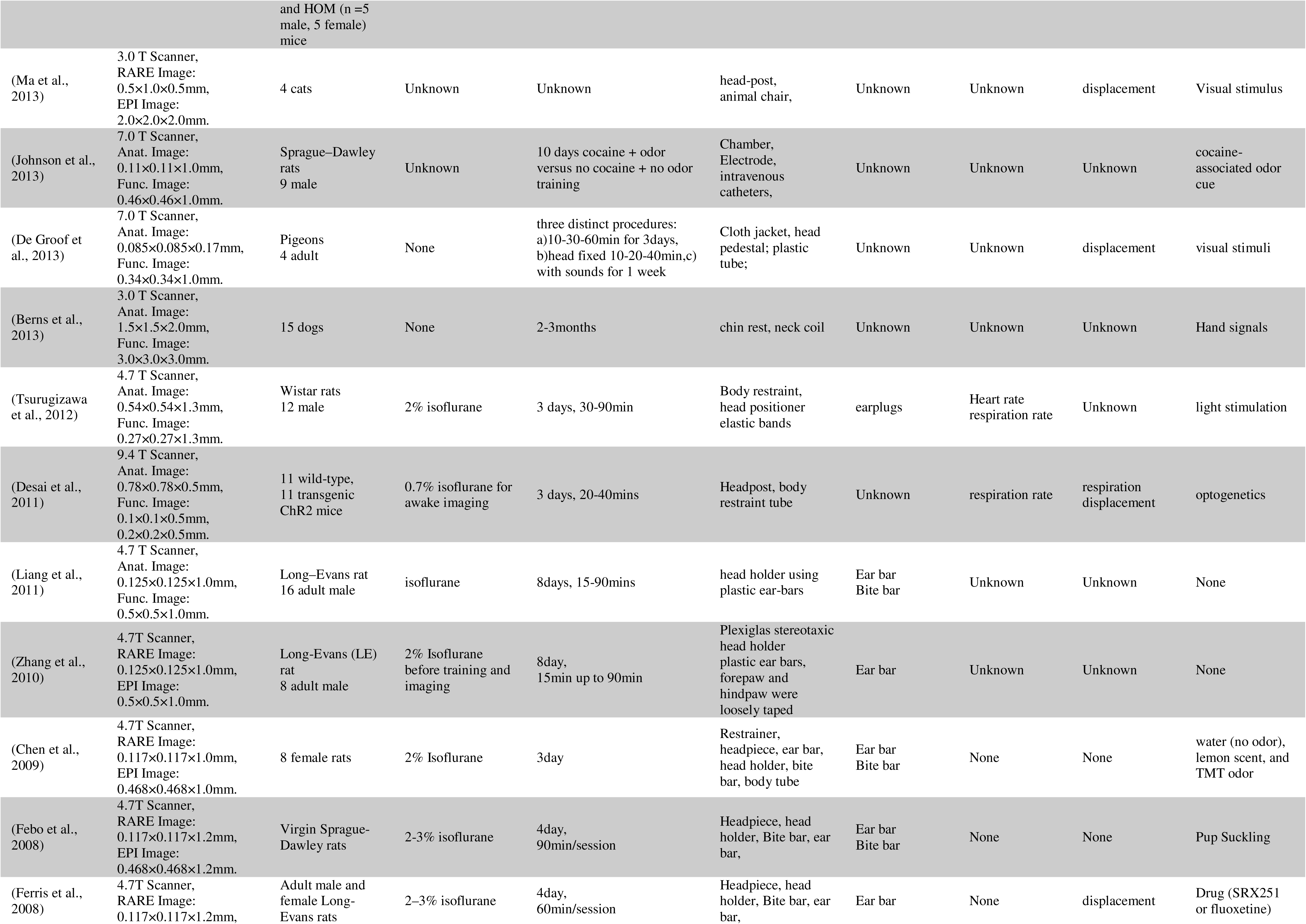
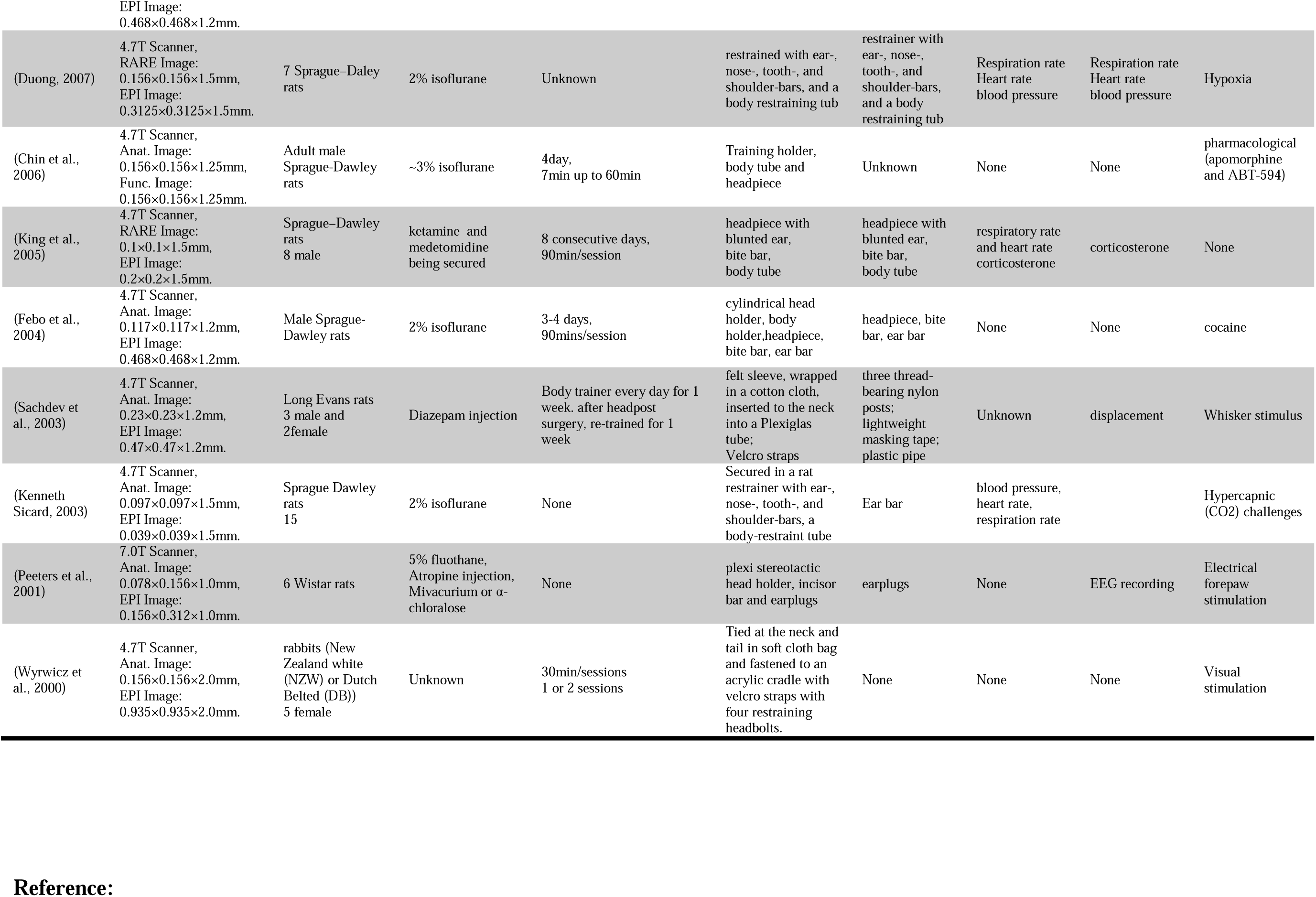
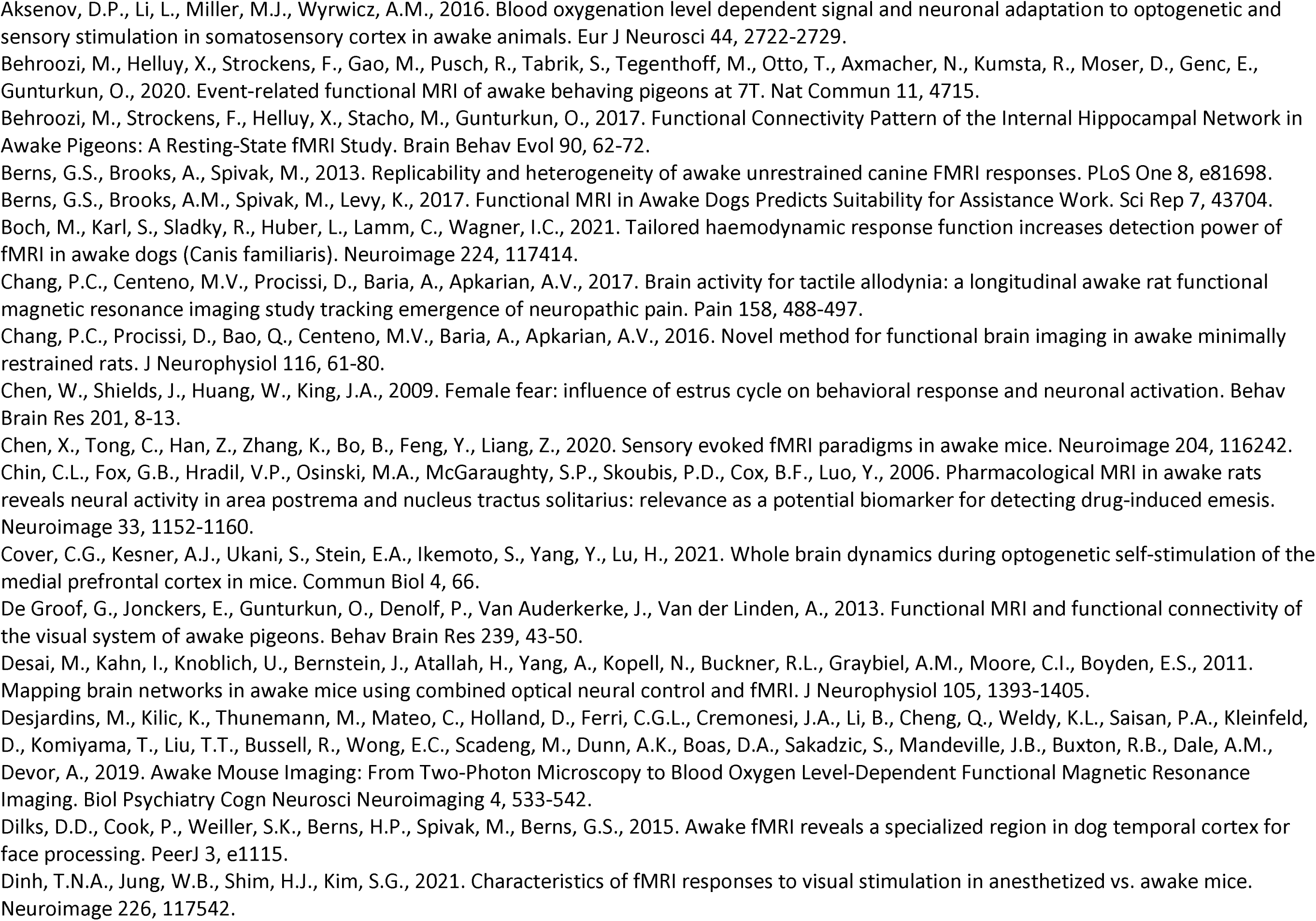
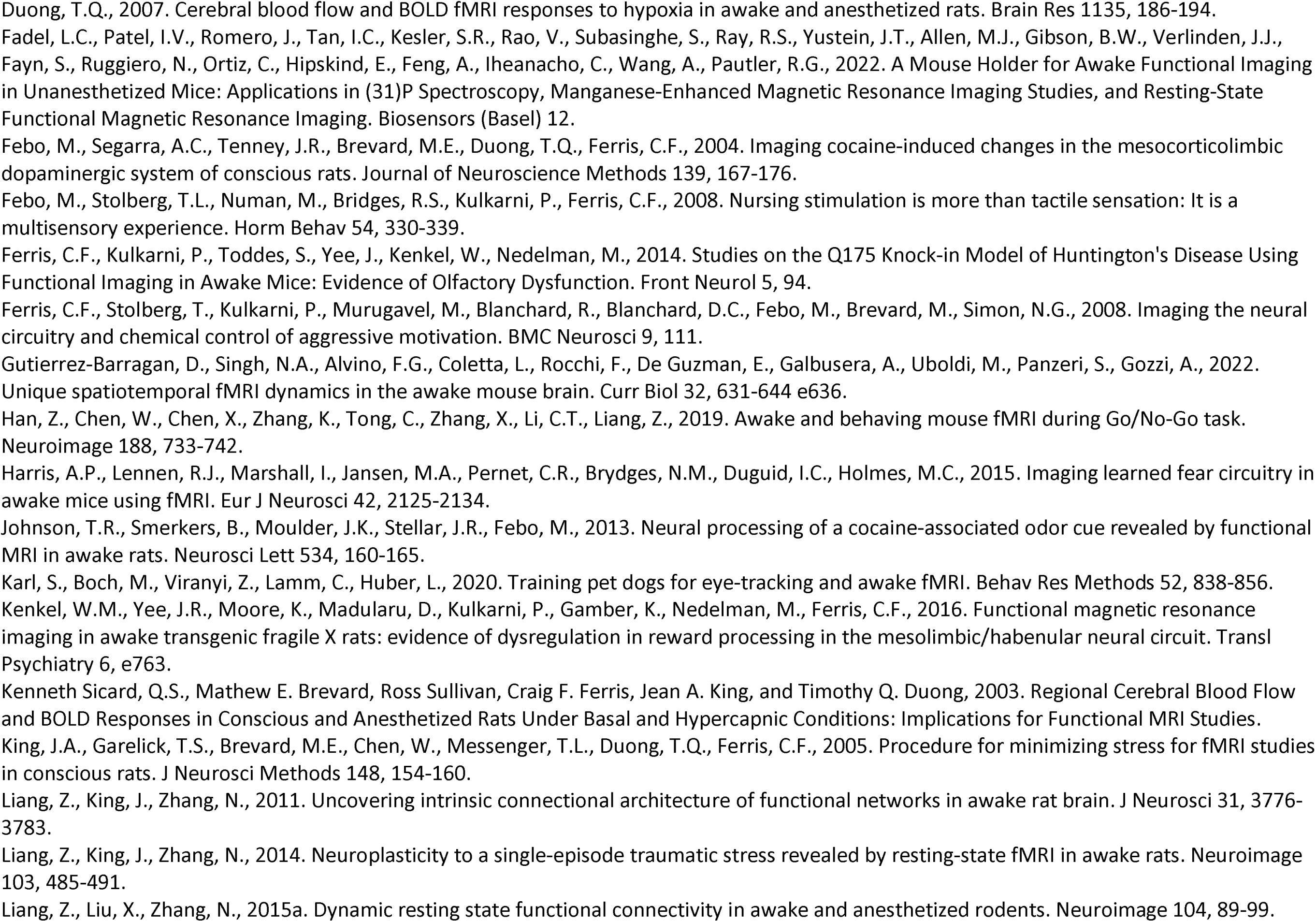
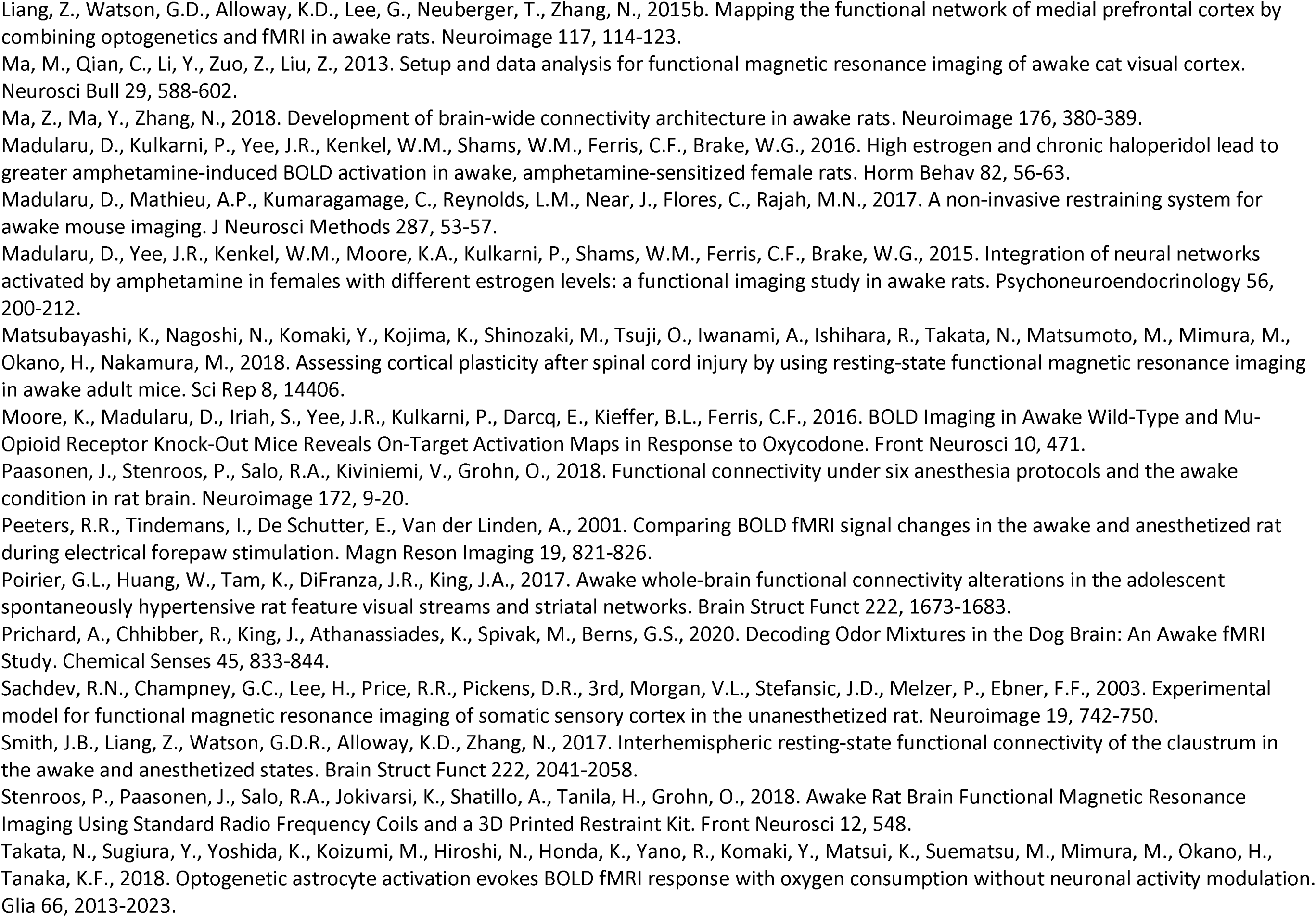
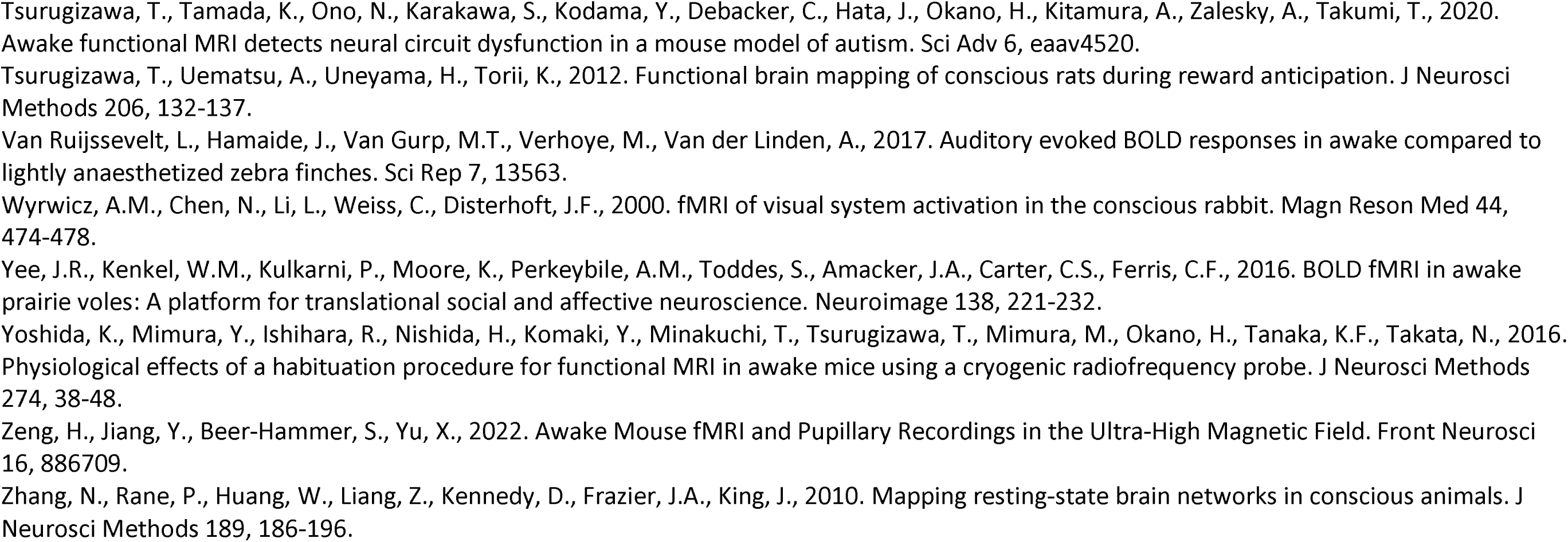

